# Cross-Species Cortical Geometry Reveals Conserved Gradients Across Primates and Human-Specific Expansion

**DOI:** 10.1101/2025.11.28.691123

**Authors:** Yifan Wei, Yufan Wang, Tingwei Wei, Xiaoting Lu, Deying Li, Chet C. Sherwood, Yu Zhang, Chen Cheng, Tianzi Jiang, Lingzhong Fan, Luqi Cheng

**Author notes:** Corresponding Author: **Luqi Cheng**, School of Life and Environmental Sciences, Guilin University of Electronic Technology, Guilin 541004, China.;., **Lingzhong Fan**, Institute of Automation, Chinese Academy of Sciences, Beijing 100190, China., **Tianzi Jiang**, Institute of Automation, Chinese Academy of Sciences, Beijing 100190, China. These authors contributed equally to this work (Yifan Wei, Yufan Wang).

## Abstract

The primate cerebral cortex, characterized by its complex structural geometry, underlies advanced cognitive functions and represents a defining feature distinguishing primates from other mammals. However, cross-species patterns of cortical geometry and the links between human cortical geometry and transcriptional architecture remain poorly understood. We developed a geometry-based cross-species cortical alignment framework to systematically investigate the similarities and differences in structural connectivity and cortical expansion characteristics among macaques, chimpanzees, and humans, and additionally explored the transcriptional underpinnings of human cortical geometry. Our analysis revealed conserved spatial patterns of cortical geometric features across species, providing the foundation for constructing a cross-species structural common space to support the alignment framework. We found that primary sensory, somatomotor, and face-selective regions exhibited high structural connectivity similarity across species, whereas prefrontal and parietal association cortices displayed significant divergence. We also identified disproportionate cortical expansion in the default mode network, with a consistent expansion trend across different evolutionary lineages in primates. Furthermore, neuroimage-transcription analysis indicated that cortical geometric features were correlated with transcriptional profiles enriched in neurodevelopmental and connectivity-related pathways. These results highlight a conserved yet hierarchically differentiated organization of the cerebral cortex in primates, providing new insights into the biological basis of human brain evolution.

## 1. Introduction

The cerebral cortex, as the highest integrative center of the central nervous system of mammals, serves as the neural substrate for advanced cognitive functions in humans, such as executive control, language, and social cognition (Friedman and Robbins 2022, Menon and D’Esposito 2022). It also represents one of the key features that distinguish primates from other mammals (Geschwind and Rakic 2013, Akula, Exposito-Alonso et al. 2023, Vanderhaeghen and Polleux 2023). Beyond functional complexity, mammalian brains displays intricate structural organization, whose three-dimensional morphology not only reflects regional functional specialization and developmental patterns, but also encodes rich biological information, including genetic ancestry (Fan, Bartsch et al. 2015, Fernández, Llinares-Benadero et al. 2016, Silva, Peyre et al. 2019, Vohryzek, Sanz-Perl et al. 2025).

Comparative analysis of neuroimaging data across species is a key approach in the study of brain evolution, encompassing a variety of methodological frameworks (Rilling 2014, Cheng, Zhang et al. 2021, Friedrich, Forkel et al. 2021). Among these, cross-species alignment plays a pivotal role, with its success largely depending on the ability to establish meaningful correspondences between cortical regions of different species (Mars, Sotiropoulos et al. 2018, Eichert, Robinson et al. 2020). Previous alignment approaches based on function and structure often assume that certain homologous regions are evolutionarily conserved and use them as anchors for interspecies comparison (Van Essen and Dierker 2007, van Heukelum, Mars et al. 2020, Xu, Nenning et al. 2020, Amano, Tanabe et al. 2025). However, definitions of homology are often grounded in human neuroanatomy and may overlook substantial interspecies differences in brain structure, regional size, and organization, thus limiting their generalizability. In addition, many of these methods require complex multimodal data and primarily focus on aligning discrete anatomical landmarks or regions of interest, without fully exploiting the global geometry and continuity of the cortical surface, which are critical for capturing complex cross-species variations in cortical topology.

Recent studies have increasingly emphasized the importance of cortical geometry, highlighting strong coupling between anatomical geometry and brain function from both functional connectivity and structural perspectives (Damoiseaux and Greicius 2009, Hilgetag, Beul et al. 2019, Robinson 2019, Meng, Yang et al. 2022, Luo, Zhang et al. 2023, Schwartz, Nenning et al. 2023, Li, Zalesky et al. 2025). Compared with complex functional networks, geometric features such as cortical shape and spatial positioning are considered more fundamental constraints of brain function, influencing the emergence and evolution of cognition (Pang, Aquino et al. 2023). These geometric properties are established during early embryonic development and exhibit a degree of evolutionary stability across species, suggesting that they form a crucial foundation for the evolution of neural processing related to a range of functions (Wedeen, Rosene et al. 2012, Valk, Xu et al. 2022). Therefore, understanding the physical geometry of the cortex and its evolutionary stability provides a fundamental basis for elucidating how structural constraints have shaped the functional and behavioral diversification of primate species.

In this study, to facilitate the comparison among species, we propose a universal cross-species alignment framework based on the conserved cortical geometric morphology. The cortical geometries were characterized by the three-dimensional coordinates from the cortical surfaces of macaques, chimpanzees, and humans, and a joint-embedding dimensionality reduction approach was applied to project the dominant geometric components into a shared low-dimensional space (Coifman and Lafon 2006, Nenning, Xu et al. 2020). This joint-embedding space preserves species-specific geometric features while capturing interspecies similarity and dissimilarity in cortical structure, providing a robust framework for cross-species comparison and a unified coordinate system for alignment. Building on this framework, we further quantified interspecies structural connectivity similarity and conducted cortical expansion analyses, thereby revealing conserved and divergent evolutionary patterns of the cortex across these three primate species. Finally, a neuroimage-transcription correlation analysis was performed to uncover transcriptional mechanisms underlying of cortical geometric features in humans.

## 2. Methods

### 2.1 Datasets

#### Human data

Forty right-handed healthy adults (aged 22–35, including 18 males) were randomly selected the Human Connectome Project (HCP) (http://www.humanconnectome.org/study/hcp-young-adult) (Van Essen, Smith et al. 2013). T1-weighted (T1w) MPRAGE images (0.7 mm isotropic resolution; 256 slices; field of view: 224 × 320; flip angle: 8°) and diffusion-weighted images (DWI) (1.25 mm isotropic resolution; 111 slices; field of view: 210 × 180; flip angle: 78°; b-values: 1000, 2000, and 3000 s/mm²) were acquired using a 3T Skyra scanner (Siemens, Erlangen, Germany) with a 32-channel head coil.

#### Chimpanzee data

Data from 27 adult chimpanzees (*Pan troglodytes*, 14 males) were obtained from the National Chimpanzee Brain Resource (http://www.chimpanzeebrain.org). Both T1-weighted (T1w) and diffusion-weighted imaging (DWI) data were collected at the Emory National Primate Research Center (ENPRC) using a 3T MRI scanner under propofol anesthesia (10 mg/kg/h), following previously described procedures (Chen, Errangi et al. 2013). All experimental protocols were approved by the ENPRC and the Emory University Institutional Animal Care and Use Committee (approval no. YER-2001206).

DWI data were acquired with a single-shot spin-echo echo-planar imaging sequence across 60 diffusion directions (b = 1000 s/mm²; repetition time = 5900 ms; echo time = 86 ms; 41 slices; 1.8 mm isotropic resolution). To correct for susceptibility-induced distortions, scans were performed using opposite phase-encoding directions (left–right). Each DWI set also included five b = 0 s/mm² images with identical imaging parameters. T1w images were acquired for each subject (218 slices; resolution: 0.7 × 0.7 × 1 mm).

#### Macaque data

Data from eight adult male rhesus macaque monkeys (*Macaca mulatta*) were obtained from TheVirtualBrain repository (https://openneuro.org/datasets/ds001875/versions/1.0.3) (Shen, Bezgin et al. 2019). All surgical and experimental procedures were approved by the Animal Use Subcommittee of the University of Western Ontario Council on Animal Care (AUP no. 2008–125) and conducted in accordance with Canadian Council of Animal Care guidelines. Surgical preparations, anesthesia protocols, and imaging procedures have been detailed previously (Shen, Bezgin et al. 2019). Imaging was performed using a 7 T Siemens MAGNETOM head scanner. For each monkey, two diffusion-weighted scans were acquired with opposite phase encoding in the superior–inferior direction at a 1 mm isotropic resolution to enable correction for susceptibility-induced distortions. For five of the animals, data were acquired using a 2D EPI diffusion protocol, while the remaining three were scanned using a multiband EPI diffusion sequence. All diffusion scans were conducted with b = 1000 s/mm², 64 diffusion directions, and 24 slices. Additionally, a 3D T1-weighted image was acquired for each subject (128 slices; 0.5 mm isotropic resolution).

### 2.2 Image preprocessing

The human T1-weighted (T1w) structural data were preprocessed using the HCP minimal preprocessing pipeline (Glasser, Sotiropoulos et al. 2013), whereas the chimpanzee and macaque T1w structural data followed the HCP’s nonhuman primate preprocessing protocols as described in prior studies (Gilissen and Hopkins 2013, Donahue, Glasser et al. 2018). In brief, the pipeline included alignment to a standard volumetric space using FSL, automated cortical surface reconstruction with FreeSurfer, and alignment to a group-averaged surface template via the Multimodal Surface Matching (MSM) algorithm (Robinson, Jbabdi et al. 2014). Human volumetric data were aligned to the Montreal Neurological Institute (MNI) space, and surface data were transformed to the fs_LR surface template. Chimpanzee data were registered to the Yerkes29 chimpanzee template, while macaque data were aligned to the Yerkes19 macaque template (Donahue, Glasser et al. 2018).

Diffusion-weighted image (DWI) preprocessing was conducted similarly across the human, chimpanzee, and macaque datasets using FSL. A diffusion tensor model was fitted for each dataset using FSL’s DTIFIT tool. Subsequently, voxel-wise estimates of fiber orientation distributions were computed using Bedpostx, with three fiber orientations modeled for the human data and two orientations for both chimpanzee and macaque data, reflecting the corresponding b-values used in acquisition.

### 2.3 Geometric Feature Extraction

To enable cross-species comparative analysis of cortical geometry, it is essential to extract structurally comparable components that are conserved across species. In this study, we developed a method to construct a shared cross-species space by leveraging geometric features of cortical surfaces, enabling the identification of corresponding components across species—termed geometric gradients. Specifically, we first extracted the three-dimensional coordinates of each vertex on the cortical midthickness surfaces for humans, chimpanzees, and macaques (**Figure 1A)**. These surface coordinate matrices were then vertically concatenated to form a joint matrix that represents the spatial layout of cortical surfaces across species (**Figure 1B**). To quantify inter-vertex relationships, we computed a cosine distance matrix based on the joint matrix. Each row of this matrix encodes the pairwise cosine distances between one vertex and all others across species, capturing the spatial proximity patterns between cortical regions (**Figure 1C**).

**Figure 1.**
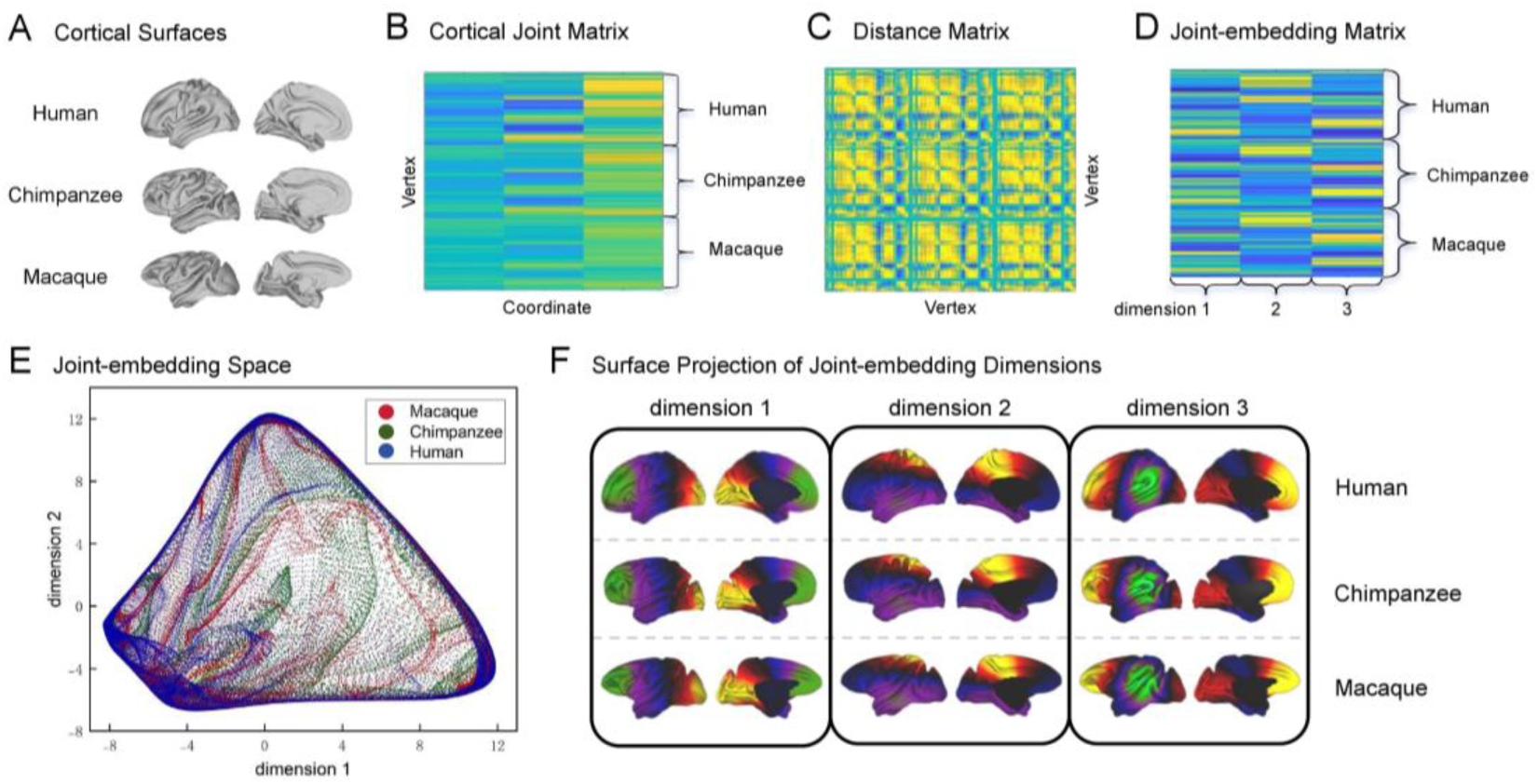
Cross-species cortical geometries captured by joint embedding. (A) Midthickness cortical surfaces of human, chimpanzee, and macaque brains. (B) Cortical joint matrix constructed by concatenating vertex-wise surface coordinates across species. (C) Cross-species distance matrix. (D) Joint-embedding matrix with the three embedding dimensions. (E) Vertex projection of the three species in the joint-embedding space. (F) The first three joint-embedding dimensions for human, chimpanzee, and macaque.

Next, we applied diffusion embedding to the distance matrix to extract a set of low-dimensional embedding components (Coifman and Lafon 2006, Nenning, Xu et al. 2020, Vos de Wael, Benkarim et al. 2020). Importantly, the resulting embedding, referred to as the joint-embedding space, maintains the same row structure as the original joint matrix: the first segment corresponds to humans, the second to chimpanzees, and the third to macaques. Each column of the embedding represents one geometric gradient dimension, capturing shared patterns of cortical geometry across species (**Figure 1D**).

Based on the three-dimensional cortical coordinates, the embedding yielded three gradient dimensions, which were retained as the foundation for constructing the cross-species comparison framework. These gradients reflect interspecies topographic similarity and enable cross-species comparison of cortical organization in a shared, low-dimensional space, improving both computational efficiency and interpretability. To further verify the cross-species consistency of cortical geometric patterns, we additionally extracted geometric features from eight primate species, including the Mangabey, Colobus Monkey, Night Monkey, Capuchin Monkey, Woolly Monkey, Saki Monkey and Galago, following the same processing pipeline to examine the similarity of geometric organization across a broader phylogenetic range.

### 2.4 Cross-Species Cortical Surface Alignment

To establish vertex-wise correspondence across human, chimpanzee, and macaque cortical surfaces, we employed Multimodal Surface Matching (MSM) (Robinson, Jbabdi et al. 2014), a spherical alignment framework that aligns surfaces based on feature similarity while minimizing distortion. The geometric gradients previously extracted were used as surface features to drive alignment across species (**Figure 2A**). The cortical surfaces of all three species were first projected onto spheres, and the first three joint-embedding gradients were selected as input features for MSM. To further enhance alignment accuracy and avoid topological misalignment, particularly in the medial wall, we incorporated two additional constraints into the MSM cost function: (1) a medial wall mask to exclude non-cortical regions, and (2) MT maps (T1w/T2w ratio) as auxiliary features, which provided complementary structural contrast sensitive to myelin content.

**Figure 2.**
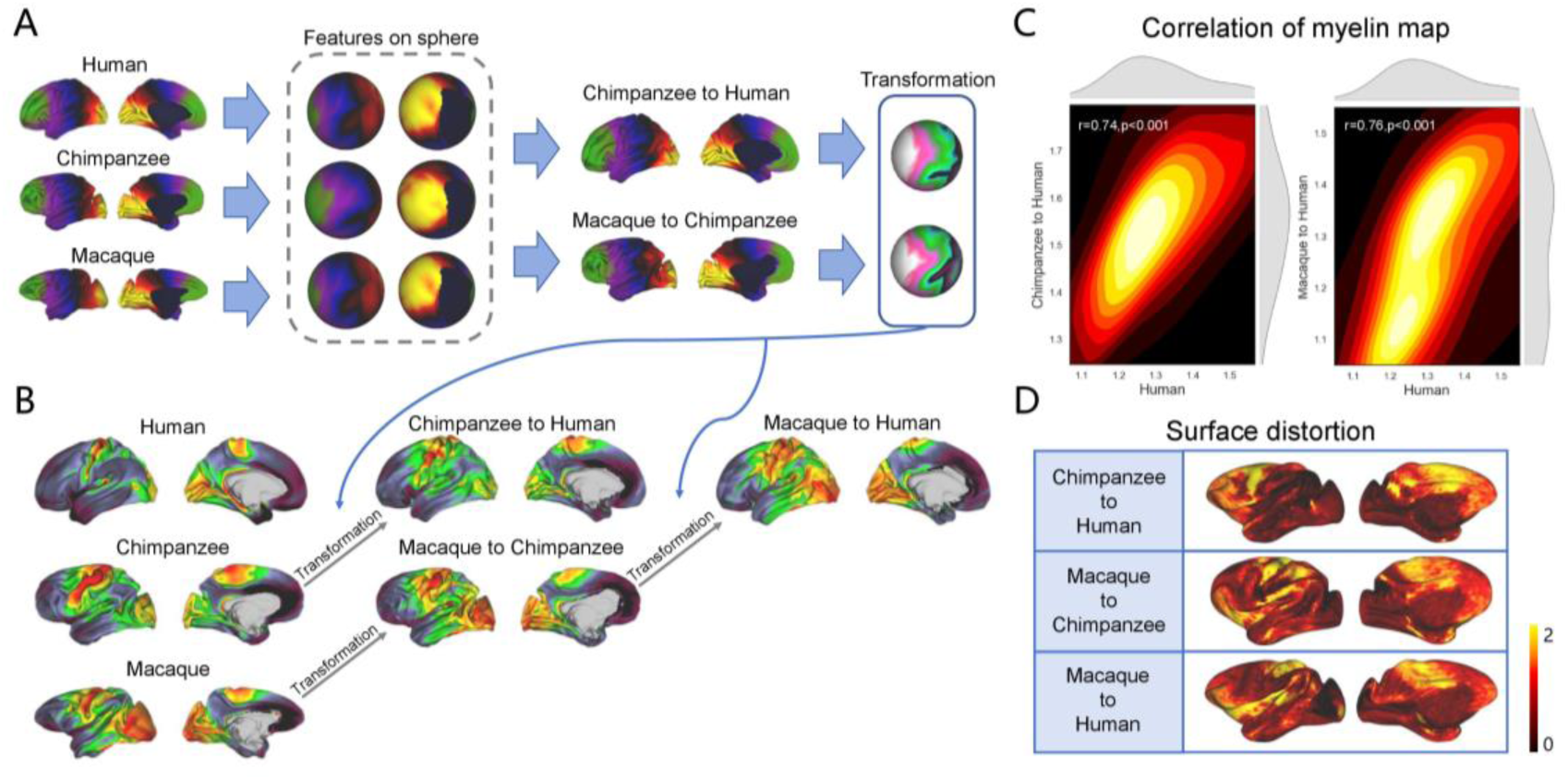
Cross-species surface alignment using geometric features. (A) Cross-species alignment framework based on geometric gradient features. (B) Transformations applied to T1w/T2w myelin maps, enabling alignment from macaque to chimpanzee and to human. (C) Correlation of cross-species myelin maps. Density plots demonstrates significant spatial correlation between the original human map and cross-species predictions. (D) Surface distortions quantifies local expansion (>1) or contraction (<1) introduced during registration.

Alignment performance was assessed by applying the computed surface deformation fields to the cortical surfaces of the source species, followed by qualitative evaluation of the transformed surfaces in the target space (**Figure 2B**). We further assessed alignment accuracy by transforming the myelin-sensitive T1w/T2w maps and quantifying their vertex-wise correlations with native maps of the target species (**Figure 2C**). In addition, we quantified surface distortion maps to evaluate the degree of geometric deformation required for alignment (**Figure 2D**). To further demonstrate the broad applicability of our alignment framework, we applied the derived cross-species transformations to the Brainnetome Atlas across macaque, chimpanzee, and human cortices (**Supplementary Figure 3**) (Fan, Li et al. 2016, Lu, Cui et al. 2024, Wang, Cheng et al. 2025).

### 2.5 Structural Connectivity Similarity Index (SCSI)

To quantitatively evaluate the local structural similarity of cortical organization across species in the joint embedding space, we developed a Structural Cortical Similarity Index (SCSI), which measures the spatial correspondence between human structural connectivity maps and those predicted from nonhuman primates (**Figure 3A**). Specifically, structural connectivity profiles (i.e. connectivity blueprints) derived from homologous white matter tracts in macaques and chimpanzees were first aligned to the human cortical template using the optimized Multimodal Surface Matching (MSM) algorithm, resulting in species-aligned predicted structural maps. (i.e., macaque predicted and chimpanzee predicted) (Robinson, Jbabdi et al. 2014, Mars, Sotiropoulos et al. 2018, Wang, Cheng et al. 2025). This surface-based alignment ensured vertex-wise correspondence between human and nonhuman cortical surfaces, allowing for spatially resolved cross-species comparison. For each vertex on the human cortical surface using the midthickness mesh, a local searchlight region with a radius of 12 mm was defined to evaluate local structural pattern similarity. Within each searchlight, cosine similarity was calculated between the structural profile of the human map and that of the predicted map derived from either macaque or chimpanzee (Mars, Sotiropoulos et al. 2018). The maximum cosine similarity within the local region was assigned as the SCSI value of the center vertex. This process generated a vertex-wise SCSI map across the human cortex, indicating the degree of local structural similarity with each nonhuman species.

**Figure 3.**
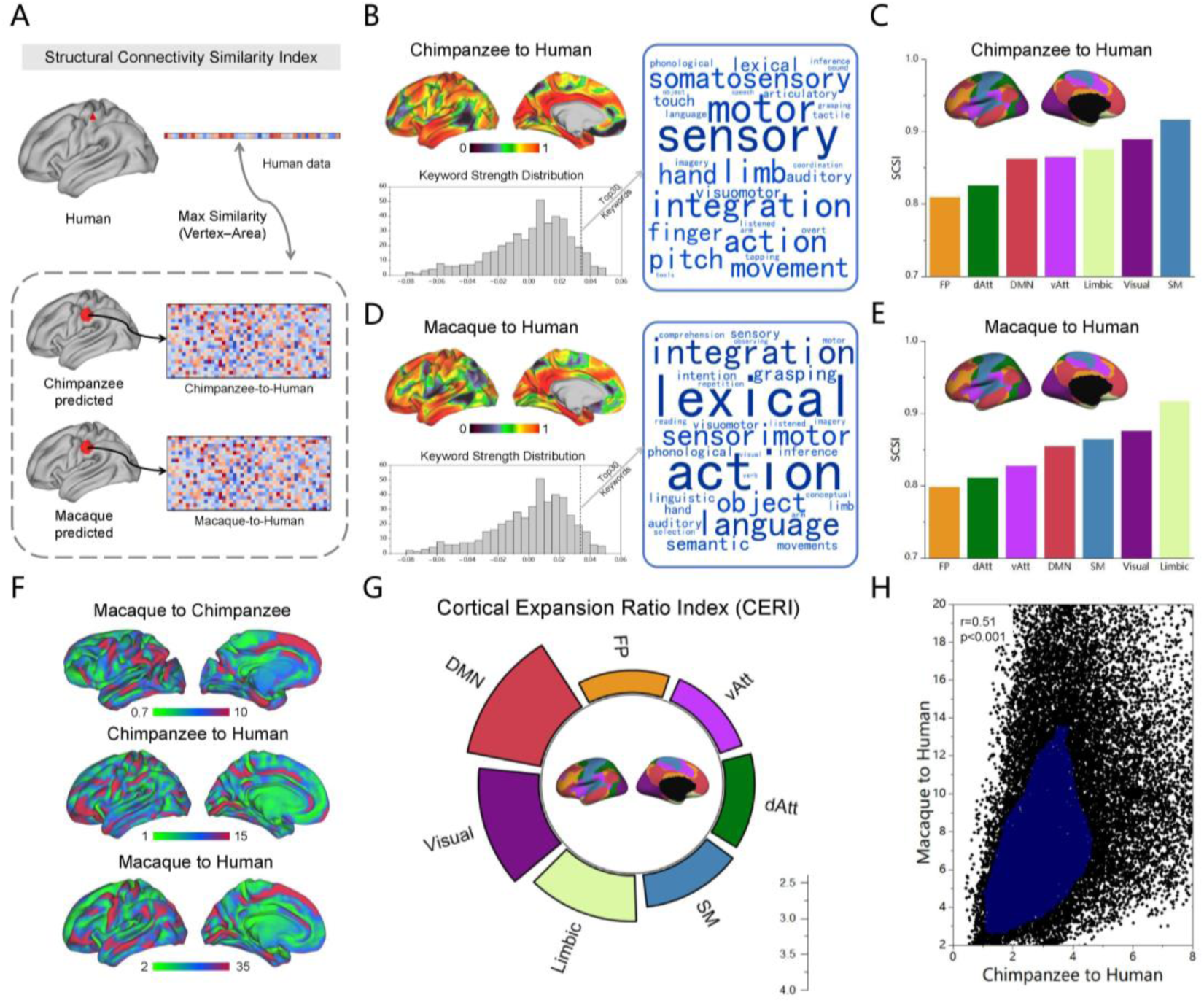
Quantification of cross-species structural connectivity similarity and cortical expansion. (A) Structural Connectivity Similarity Index (SCSI). (B) Chimpanzee-to-human SCSI map with meta-analytic decoding. (C) Chimpanzee-to-human SCSI across seven functional networks defined by Yeo et al. (2011). (D) Macaque-to-human SCSI map with meta-analytic decoding. (E) Macaque-to-human SCSI across canonical functional networks. (F) Cortical expansion maps for macaque-to-chimpanzee, chimpanzee-to-human, and macaque-to-human comparisons. (G) Cortical Expansion Ratio Index (CERI). (H) Vertex-wise correlation between chimpanzee-to-human and macaque-to-human expansion maps.

To quantify the relationship between cross-species structural similarity and cognitive function, we performed meta-analytic functional decoding of the SCSI maps using the NeuroSynth database (Yarkoni, Poldrack et al. 2011). The SCSI maps were correlated with meta-analytic functional activation maps related to distinct cognitive and behavioral terms, and the top 20 most strongly correlated terms were retained for further interpretation.

### 2.6 Cortical Expansion

To quantify cortical expansion during primate evolution, we computed a vertex-wise expansion index across species based on point-to-point correspondences derived from the alignment method. Specifically, vertex-wise surface area was estimated for all individuals on the 32k standard surface mesh and averaged within each species to generate species-specific mean surface area maps. The macaque and chimpanzee maps were then aligned to the human target space using MSM-derived alignment spheres (Robinson, Jbabdi et al. 2014), and the macaque maps were also aligned to chimpanzee space, thereby establishing vertex-wise correspondences across species. Based on these correspondences, three expansion maps were generated: macaque-to-chimpanzee, chimpanzee-to-human, and macaque-to-human (**Figure 3F**). For the chimpanzee-to-human and macaque-to-human comparisons, expansion at each vertex was defined as the ratio of human surface area to the corresponding nonhuman primate surface area after alignment. In contrast, macaque-to-chimpanzee expansion was computed directly in chimpanzee space by aligning macaque surfaces to chimpanzees and calculating vertex-wise area ratios.

Beyond the vertex-wise expansion maps, we further quantified the relative contributions of different evolutionary stages by defining the Cortical Expansion Ratio Index (CERI). CERI was computed as the ratio of macaque-to-human expansion to chimpanzee-to-human expansion at each vertex, thereby highlighting regions that exhibit disproportionate expansion in earlier (macaque-to-chimpanzee) versus later (chimpanzee-to-human) branch points in primate evolution (**Figure 3G**).

### 2.7 Estimation of regional gene expressions

To investigate spatial variations in gene expression across the human cortex, we utilized microarray transcriptional data from the Allen Human Brain Atlas (Hawrylycz, Lein et al. 2012, Sunkin, Ng et al. 2012), which includes genome-wide expression profiles obtained from 3,702 spatially distributed tissue samples across six adult postmortem brains (male/female = 5/1; age = 42.5 ± 13.4 years). Due to the limited and inconsistent coverage of the right hemisphere (only two donors), only data from the left hemisphere were retained for subsequent analysis to minimize sampling bias. Data preprocessing followed the standardized pipeline proposed by Arnatkeviciute et al. (2019) and was implemented using the Python-based toolbox abagen (Arnatkeviciūtė, Fulcher et al. 2019, Markello, Arnatkeviciute et al. 2021).

The processing steps included: (1) re-annotation of probe-to-gene mappings using the Re-annotator toolkit; (2) exclusion of probes with low expression intensity (i.e., below background noise in >50% of samples); (3) selection of the most reliable probe per gene based on maximal correlation with RNA-seq data; (4) mapping AHBA samples to regions defined by the Human Brainnetome Atlas (Fan, Li et al. 2016); (5) normalization of expression values across participants using a scaled robust sigmoid function to account for inter-individual variability; (6) filtering of genes based on differential stability to retain only genes with consistent spatial expression patterns across donors. This processing resulted in a regional gene expression matrix consisting of 15,633 genes across 105 cortical regions in the left hemisphere, which was used for subsequent spatial correlation analysis.

### 2.8 Neuroimaging–transcriptomic association analysis of cortical geometry

To investigate the transcriptomic correlates of regional variation in brain phenotypes, we performed partial least squares (PLS) regression (Krishnan, Williams et al. 2011, Abdi and Williams 2012). We used a gene expression matrix comprising 15,633 genes across 105 cortical regions (**Figure 4A**), derived based on the Human Brainnetome Atlas (Fan, Li et al. 2016), as the independent variable X, and the region-wise brain geometry as the dependent variable Y. Partial least squares (PLS) regression was conducted to identify gene expression patterns associated with human cortical geometry. Only the first component (PLS1) was retained for further analysis, as it explained the greatest variance. The statistical significance of PLS1 was assessed using a spherical permutation test (n = 5000), yielding an empirical p-value (*p* < 0.001) (Váša, Seidlitz et al. 2018).

**Figure 4.**
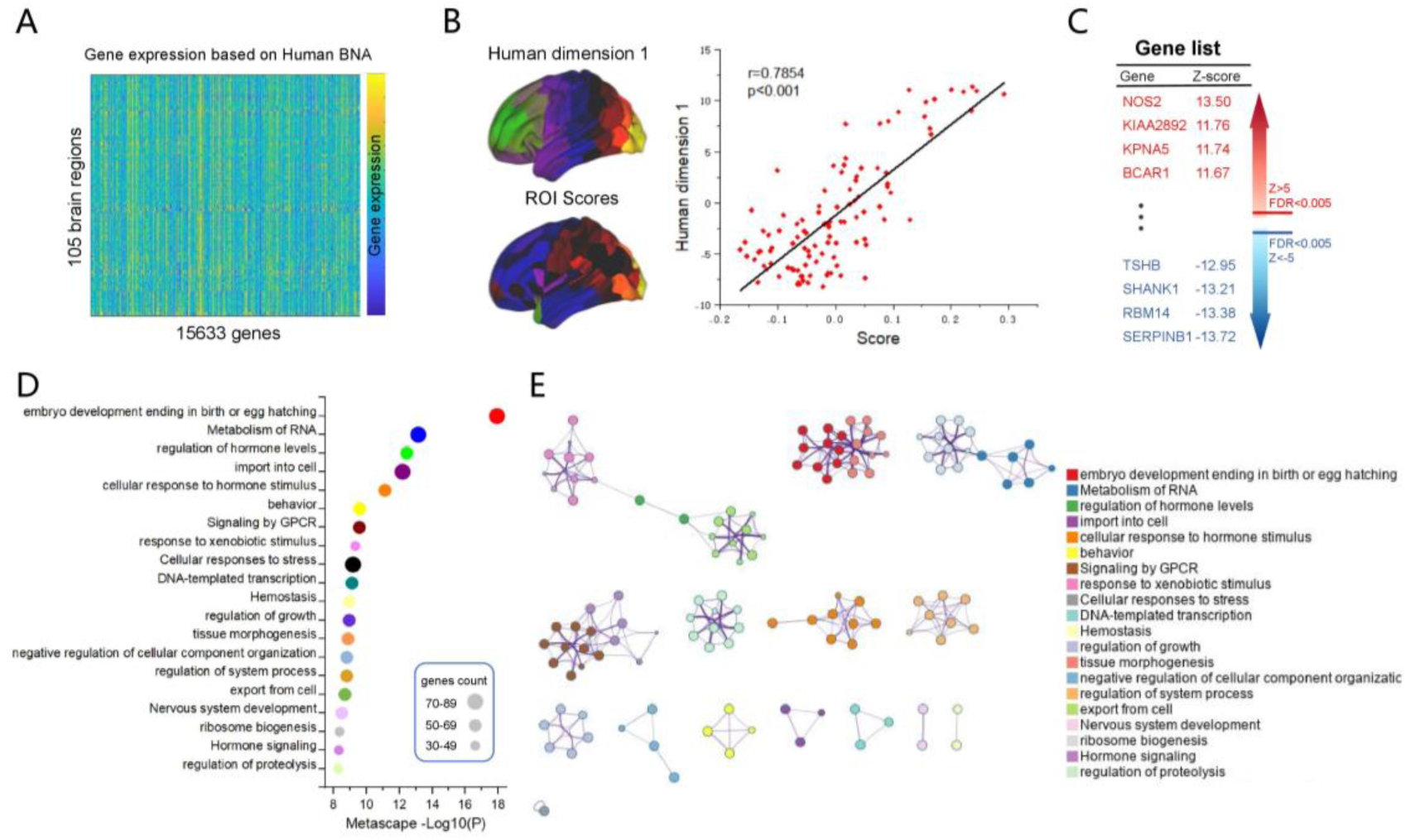
Transcriptomic correlates of human cortical geometry. (A) Regional gene expression matrix derived from the Allen Human Brain Atlas (AHBA) and parcellated based on the Human Brainnetome Atlas. (B) Cortical maps of human dimension 1 and regional PLS1 gene expression weighted values. Right panel shows a significant correlation between PLS1 scores and regional values of dimension 1. (C) The PLS1 gene expression weights (|Z| > 5 and FDR < 0.005). (D) Functional enrichment analysis of PLS1+ genes using Metascape, revealing overrepresented Gene Ontology (GO) biological processes. Circle size represents the number of genes per term; color denotes cluster identity. (E) Network visualization of enriched GO terms, where nodes represent individual terms and terms grouped by color reflect distinct functional modules.

To evaluate the robustness of gene weights in PLS1, bootstrapping (5,000 iterations) was conducted by resampling brain regions with replacement and re-computing PLS regression. For each gene, a Z-score was calculated by dividing the original PLS1 weight by the standard deviation of the bootstrapped weights (Morgan, Seidlitz et al. 2019). Subsequently, false discovery rate (FDR) correction was applied to the p-values converted from Z-scores. Genes with |Z| > 5 and FDR < 0.005 were retained for further analysis. Genes with significantly positive Z-scores were denoted as PLS1+, while those with significantly negative Z-scores were denoted as PLS1−(**Figure 4C**).

To interpret the biological significance of genes positively associated with the brain geometry, functional enrichment analysis was performed using Metascape (Zhou, Zhou et al. 2019), an online resource that integrates over 40 independent knowledgebases for gene annotation and pathway analysis. The PLS1+ gene list (Z > 5 and FDR < 0.005) derived from PLS regression was submitted to Metascape to identify overrepresented biological functions. Gene Ontology (GO) Biological Process (Ashburner, Ball et al. 2000) and Kyoto Encyclopedia of Genes and Genomes (KEGG) pathway databases were selected to explore the molecular pathways enriched among PLS1+ genes. The significance threshold for all enriched pathways was set at FDR-corrected *p* < 0.05. This analysis enabled identification of biological processes and signaling pathways potentially underlying the transcriptomic architecture of regional brain alterations.

## 3. Results

### 3.1 Construction of a Cross-Species Structural Common Space

We first applied diffusion embedding to the distance matrix of cortical coordinates to characterize the geometric gradients for macaques, chimpanzees, and humans. Given the three-dimensional nature of the cortical coordinates, three geometric feature gradients were derived for macaques, chimpanzees, and humans (**Figure 1D**). This approach effectively preserved cortical geometric information, with each embedding dimension representing a common feature along its respective axis. We then projected each vertex of the three species into the joint-embedding space defined by the embedding dimensions. Notably, the embeddings of humans, chimpanzees, and macaques showed strong correspondence within the shared coordinate system (**Figure 1E**).

To further interpret these embedding dimensions, we examined their spatial organizations (**Figure 1F**). Dimension 1 followed an anterior-posterior gradient, spanning from occipital and temporal sensory areas to frontal association regions. Dimension 2 displayed a dorsal-ventral organization, while dimension 3 captured variation along the medial–lateral axis. These gradients exhibited highly similar spatial distributions and conserved patterns across the three species, thereby providing a cross-species structural common space for subsequent cross-species alignment. Meanwhile, we extended the analysis to additional primate species **(Supplementary Figure 2) (Bryant, Ardesch et al. 2021)**. The resulting patterns closely resembled those observed in humans, chimpanzees, and macaques, indicating a broadly conserved geometric organization across primate evolution.

Additionally, we labeled homologous primary cortical regions, including the primary motor cortex (BA4), primary visual cortex (V1), and primary auditory cortex (A1) (Glasser, Coalson et al. 2016, Donahue, Glasser et al. 2018, Paxinos, Petrides et al. 2023), across the three species. The results showed that the primary cortical regions were consistently located in close proximity within the joint-embedding space (**Supplementary Figure 1**).

### 3.2 Cross-Species Alignment Guided by Geometric Features

The highly similar and conserved gradient patterns across the three species provided the basis for constructing the cross-species alignment framework (**Figure 2A**). For each species, the three gradients were selected as input geometric features. We first aligned macaque features to the target chimpanzee space, yielding a macaque-to-chimpanzee transformation that captured cortical surface deformations. Next, chimpanzee features were aligned to the target human space, generating a chimpanzee-to-human transformation. For macaque-to-human alignment, this framework employed chimpanzees as an intermediate reference.

To assess alignment performance, we compared the human myelin (T1w/T2w) map with the predicted myelin maps from macaques and chimpanzees (Glasser and Van Essen 2011). The results showed strong Spearman correlations between the empirical human myelin map and the predicted maps (chimpanzee-to-human: *r* = 0.74; macaque-to-human: *r* = 0.76; both *p_spin_* < 0.001) (**Figure 2C**). To further evaluate geometric distortion introduced by the alignment process, we analyzed mesh distortion across species (**Figure 2D**). The deformation for macaque-to-human alignment showed great distortion, particularly in temporal and prefrontal regions, whereas macaque-to-chimpanzee deformation was relatively minor. In addition, the successfull alignment of the Brainnetome Atlas across humans, macaques, and chimpanzees(**Supplementary Figure 3**) suggested the robustness and generalizability of the proposed framework.

### 3.3 Structural Connectivity Similarity Index and Cortical Expansion across Primates

Building upon the cross-species alignment framework, we next quantified similarities and differences of cortical structural connectivity (i.e. connectivity blueprints) between nonhuman primates and humans by introducing the Structural Connectivity Similarity Index (SCSI) (**Figure 3A**). This index measures the highest structural connectivity similarity between each vertex in the human cortex and regions in the cortex of the two nonhuman primates (**Figure 3B**, upper left of 3D).

For the chimpanzee-to-human comparison, higher SCSI values were observed in primary visual cortex (V1), auditory cortex, somatomotor cortex, and face-selective areas such as the fusiform face complex (FFC), while lower connectivity similarity was found in the prefrontal cortex, parietal association cortex, and posterior medial regions. Further analysis across seven canonical functional networks revealed that the somatomotor (SM) and visual networks exhibited the highest connectivity similarity, whereas the frontoparietal (FP) and dorsal attention (dAtt) networks showed the lowest similarity (**Figure 3C**), suggesting potential evolutionary remodeling of these higher-order networks in humans (Thomas Yeo, Krienen et al. 2011). Meta-analytic decoding using the NeuroSynth database indicated that regions with higher SCSI values were significantly associated with functional terms such as “sensory,” “motor,” “limb,” “integration,” and “action”.

For the macaque-to-human comparison, the overall SCSI distribution was slightly lower than that of chimpanzees, though primary sensory regions still showed relatively high similarity. Network-level analysis demonstrated that the limbic, visual, and somatomotor networks remained structurally conserved, while frontoparietal (FP) and dorsal attention (dAtt) networks continued to exhibit the lowest cross-species similarity. Similarly, NeuroSynth-based functional decoding revealed enrichment of functional topics such as “lexical,” “action,” “integration,” “language,” “sensor,” and “motor” (**Figure 3E**) (Yarkoni, Poldrack et al. 2011).

To systematically compare the relative expansion across functional networks, we proposed the Cortical Expansion Ratio Index (CERI), defined as the ratio of macaque-to-human expansion to chimpanzee-to-human expansion for each vertex. This index reflects the dominant phase of cortical expansion for a given region. Analysis across the seven functional networks (Thomas Yeo, Krienen et al. 2011) revealed that the default mode network (DMN), visual, and limbic systems exhibited the highest CERI values, indicating preferential expansion during earlier evolutionary transitions, potentially linked to the emergence of human-specific cognitive functions. Finally, we assessed the consistency between the macaque-to-human and chimpanzee-to-human cortical expansion patterns (**Figure 3H**). Correlation analysis showed a significant positive relationship (*r* = 0.51, *p_spin_* < 0.001), suggesting that despite distinct evolutionary trajectories, certain regions exhibit conserved expansion trends across primates.

### 3.4 Gene Expression and Functional Enrichment Associated with Cortical Geometry

To investigate the transcriptional underpinnings of cortical geometry, we employed partial least squares (PLS) regression to identify gene expression patterns associated with cortical geometric features in the human brain. The first component of the PLS (PLS1) represents the linear combination of gene expression values that most strongly correlates with cortical geometry, explaining over 60% of the variance in the phenotype (*p* < 0.001). Accordingly, PLS1 was selected for subsequent analysis.

Notably, we observed a significant positive correlation between the first dimension of cortical geometry and the PLS1 gene expression scores (Pearson’s r = 0.7854, *p* < 0.001; **Figure 4B**, right panel). Following the identification of PLS1 as the principal component, a bootstrapping procedure was performed to estimate the variability of each gene’s PLS1 weight. Z-scores were calculated by dividing each gene’s PLS1 weight by the standard deviation of its bootstrapped weights. Genes were ranked by their normalized PLS1 weights. This analysis identified 1,339 positively weighted genes (PLS1+, Z > 5) and 2,011 negatively weighted genes (PLS1−, Z <−5), all passing the FDR-corrected threshold of *p* < 0.05.

To gain insight into the functional roles of these genes, we performed Gene Ontology (GO) biological process enrichment analysis for both the PLS1+ and PLS1− gene sets. The PLS1+ genes were significantly enriched in GO terms such as “embryo development ending in birth or egg hatching,” “regulation of hormone levels,” “import into cell,” “cellular response to hormone stimulus,” and “behavior” (**Figure 4D–E**). Meanwhile, PLS1− genes were enriched in processes such as “cellular response to cytokine stimulus,” “cell population proliferation,” “cell activation,” “tissue morphogenesis,” “muscle structure development,” and “regulation of anatomical structure size” (**Supplementary Figure 4**).

## 4. Discussion

In this study, we developed a cross-species framework that enables robust alignment using a shared low-dimensional common space derived from the cortical geometry of macaques, chimpanzees, and humans. Using this framework, we identified interspecies variations in structural connectivity and patterns of cortical expansion in primates, and further conducted gene-level analyses to uncover the genetic underpinnings of human cortical geometry. This study provides important insights into evolutionary neuroscience, highlighting the association between cortical geometry and evolution, as well as its potential role in shaping human functional capacities.

The extracted geometric gradients revealed similar patterns across species within the same embedding dimensions (**Figure 1F**). In particular, the dominant gradient (dimension 1) exhibited a pronounced posterior-to-anterior organizational pattern, spanning from sensory areas in the occipital and temporal lobes to higher-order association regions in the frontal lobe. Interestingly, this pattern is highly consistent with the posterior-to-anterior gradients reported in previous studies (Valk, Xu et al. 2020, Li, Wang et al. 2025). This axis reflects a hierarchical transition from sensory processing to abstract cognitive functions and is known to be strongly influenced by genetic factors. Notably, similar anterior–posterior gradients have also been observed in the cortices of nonhuman primates (Cahalane, Charvet et al. 2012, Riley, Qi et al. 2018), suggesting that this organizational principle may represent an evolutionarily conserved structural framework. Such a structure could offer anatomical underpinnings for cross-species differences in cognition and shed light on the evolutionary conservation and specialization of different cortical regions. Moreover, by examining additional primate species, we further confirmed that the identified geometric organization represents a broadly conserved and evolutionarily general principle across the primate lineage, underscoring the universality of the observed cortical geometry (**Supplementary Figure 2**). This analytical framework can be readily adapted to include additional species in future studies. Our geometry-guided alignment framework demonstrated high accuracy, achieving a correlation of 0.76 in macaque-to-human myelin map prediction, demonstrating high effectiveness and robustness (Eichert, Robinson et al. 2020, Xu, Nenning et al. 2020). Furthermore, when applied to the Brainnetome Atlas, the framework produced reliable cross-species parcellation correspondences (**Supplementary Figure 3**) (Fan, Li et al. 2016, Lu, Cui et al. 2024, Wang, Cheng et al. 2025), indicating strong robustness and broad applicability.

We introduced the Structural Connectivity Similarity Index (SCSI) to quantify cross-species similarity in cortical structural connectivity. The results showed that primary cortical regions exhibited higher SCSI values. For instance, the high SCSI observed in the primary visual cortex suggests that visual processing mechanisms are highly conserved between humans and nonhuman primates, supporting the notion of structural connectivity conservation in primary cortices. In contrast, prefrontal regions such as the dorsolateral prefrontal cortex (dlPFC), which are associated with executive and social cognition in humans, showed low SCSI values—implying that these regions may have undergone significant functional reorganization during human evolution (Carlén 2017, Ma, Skarica et al. 2022). Notably, the overall SCSI values from macaques to humans were lower than those from chimpanzees, especially in high-order cognitive regions such as the prefrontal cortex, although high similarity remained in primary sensory areas. This is consistent with previous studies suggesting that macaques, being more evolutionarily distant from humans than chimpanzees, display greater divergence of structural connectivity (Croxson, Forkel et al. 2018). At the same time, the findings also underscore that while high-order cognitive areas have diverged, the organization of lower-order sensory regions has remained relatively conserved (Song, Kennedy et al. 2014, Rushworth, Goulas et al. 2019).

We further conducted a meta-analysis of the SCSI distribution using the NeuroSynth database to explore the behavioral relevance of structural connectivity similarity (Yarkoni, Poldrack et al. 2011). For chimpanzee-to-human comparisons, SCSI values were most associated with terms such as “sensory” and “motor” suggesting that these basic functions likely emerged earlier in evolution. Interestingly, in macaque-to-human comparisons, SCSI was not only associated with lower-level functions like “action” and “sensor,” but also with higher-level cognitive terms such as “lexical,” “language,” and “integration”. Given that macaques are more distantly related to humans than chimpanzees and follow a distinct evolutionary trajectory (Nakahara, Hayashi et al. 2002, Patel, Yang et al. 2015), this result suggests that the geometric basis for human higher cognitive functions may have emerged earlier in evolution, and that macaque-human cortical differences provide further insight into the origins of such capacities.

In the analysis of cross-species cortical expansion, consistent with the findings above, most primary cortical areas, such as the primary motor cortex, exhibited minimal expansion, suggesting a high degree of structural conservation throughout evolution. In contrast, the prefrontal and parietal cortices showed substantial expansion, particularly along the macaque-to-human axis, where expansion ratios exceeded 20-fold. Moreover, in the network-based analysis of the Cortical Expansion Ratio Index (CERI) derived from seven functional networks (Thomas Yeo, Krienen et al. 2011), the default mode network (DMN) exhibited the highest index values. This suggests that the DMN may have underwent the most significant expansion along the macaque–chimpanzee–human trajectory, and that such expansion likely occurred after the divergence of humans and chimpanzees, potentially contributing to human-specific cognitive enhancements. Notably, the expansion magnitudes from chimpanzee to human were significantly correlated with those from macaque to human (**Figure 3H**), suggesting a consistent expansion trend across different evolutionary stages in primates (Chaplin, Yu et al. 2013).

Gene expression associated with cortical geometric features represents a key molecular correlate and was a major focus of our study. We found that cortical geometry in the human brain was highly correlated with regional gene expression levels (**Figure 4B**), suggesting that cortical morphology may be influenced by specific transcriptional regulation patterns. This coupling provides crucial insight into the molecular basis of cortical evolution. Gene enrichment analysis further revealed that the most prominent functional terms in the PLS1+ gene set included “embryo development ending in birth or egg hatching” and “metabolism of RNA.” Given that the brain is among the earliest organs to develop during embryogenesis, these regulatory mechanisms are likely essential for shaping cortical geometry during early developmental stages (Ball, Seidlitz et al. 2020, Vasung, Zhao et al. 2021). In addition, RNA metabolism plays a key role in neuronal maintenance and function, involving transcription, splicing, modification, and degradation. These processes are critical for neuronal subtype differentiation, synaptic plasticity, and local translation, indicating that the genes influencing cortical geometry function both during embryonic development and in the maintenance of adult brain function (Grasby, Jahanshad et al. 2020, Hofer, Roshchupkin et al. 2020, Makowski, van der Meer et al. 2022, Warrier, Stauffer et al. 2023).

Nonetheless, our study has several limitations. Although cortical surface data from additional primate species were included to verify the cross-species consistency of geometric features, these datasets lacked multimodal information such as structural connectivity data. Therefore, the main analyses were restricted to humans, chimpanzees, and macaques, which provide well-characterized multimodal datasets for establishing and validating the framework. This limitation may constrain a comprehensive understanding of how cortical geometry interacts with other modalities across a broader evolutionary spectrum. Future studies incorporating multimodal data from a wider range of species will help to further elucidate the evolutionary principles shaping cortical organization. In addition, owing to the lack of gene expression data for nonhuman primates, our transcriptomic analysis was restricted to the human brain, preventing a systematic cross-species comparison of gene regulation in expanded cortical areas. Additionally, a general limitation in this field is that cortical gene expression data are derived from only six postmortem healthy adult human brains (mean age = 43 years), which limits the ability to draw definitive conclusions about the stability of gene expression across the human population. Future studies will seek to obtain transcriptomic data from additional species to enable comparative analyses across species and provide deeper insights into the biological basis of cortical geometry. Moreover, our study focused primarily on cortical regions and did not include subcortical nuclei. Future work integrating subcortical structures could optimize and extend the current framework, providing a more comprehensive understanding of brain evolution.

In summary, this study developed a high-precision cross-species cortical alignment framework based on the cortical geometry of macaques, chimpanzees, and humans. Our results highlight that primate cortical structure exhibits conserved yet hierarchically differentiated evolutionary trajectories. Across geometry, function, and gene expression, we demonstrated both shared principles and species-specific specializations between nonhuman primates and humans. Altogether, this work provides a unified geometry-based perspective for cross-species neuroimaging analysis and offers novel insights into the structural and functional evolution of the primate brain.

## Acknowledgments

This work was supported by STI2030-Major Projects (Grant No. 2021ZD0200203), Guangxi Natural Science Foundation (Grant No. 2024GXNSFBA010212), Natural Science Foundation of China (Grant Nos. 82202253, 62336007, 62201519), the China Postdoctoral Science Foundation (2022M722915), Guangxi Science and Technology Base and Talent Special Project (Grant No. AD22035125), the Basic Scientific Research Ability Improvement Project for Young and Mid-Career Teachers in Universities of Guangxi (Grant No. 2022KY0180), Central Guided Local Science and Technology Development Project (Grant No. YDZJSX2024C004), and Guangxi Academy of Artificial Intelligence. Data were provided in part by the National Chimpanzee Brain Resource (supported by NIH NS092988, NIH HG011641, NIH AG067419, NIH AG087945, NSF EF-2021785, and NSF DRL-2219759), Human Connectome Project from WU-Minn Consortium (Principal Investigators: David Van Essen and Kamil Ugurbil; 1U54MH091657) funded by the 16 NIH Institutes and Centers that support the NIH Blueprint for Neuroscience Research and by the McDonnell Center for Systems Neuroscience at Washington University.

## Data availability

The human data are available from the Human Connectome Project (https://www.humanconnectome.org). The chimpanzee data are available at the National Chimpanzee Brain Resource (http://www.chimpanzeebrain.org). The TVB macaque data are available at https://doi.org/10.18112/openneuro.ds001875.v1.0.3. All data generated or analyzed during this study are included in the manuscript and supporting files. All cortical maps generated in this study are openly available at https://github.com/ChengLabX/cortical_geometry.

