## Supplementary Figure 1-4 for "Cross-Species Cortical Geometry Reveals Conserved Gradients Across Primates and Human-Specific Expansion"

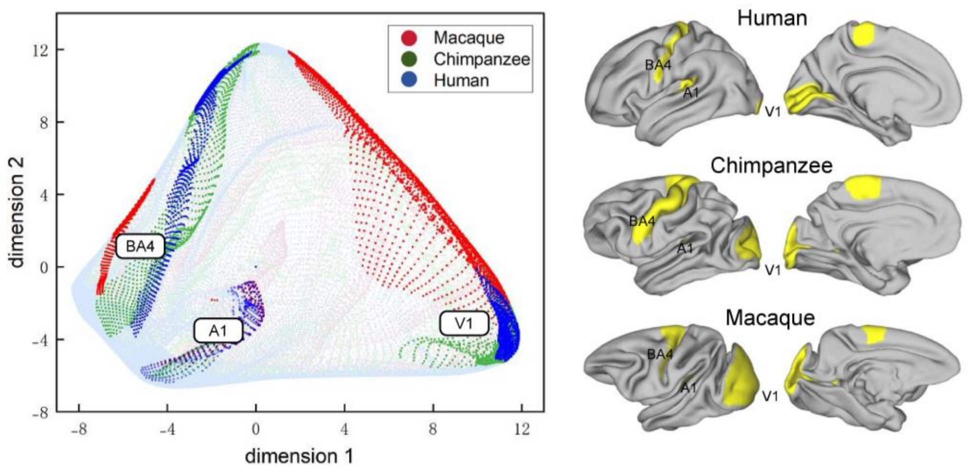


**Supplementary Figure 1.** Cross-species correspondence of primary cortical areas in the joint-embedding space. Projection of cortical vertices from human, chimpanzee, and macaque into the joint-embedding space reveals close spatial proximity of homologous primary cortical regions, including primary motor cortex (BA4), primary visual cortex (V1), and primary auditory cortex (A1) across species.

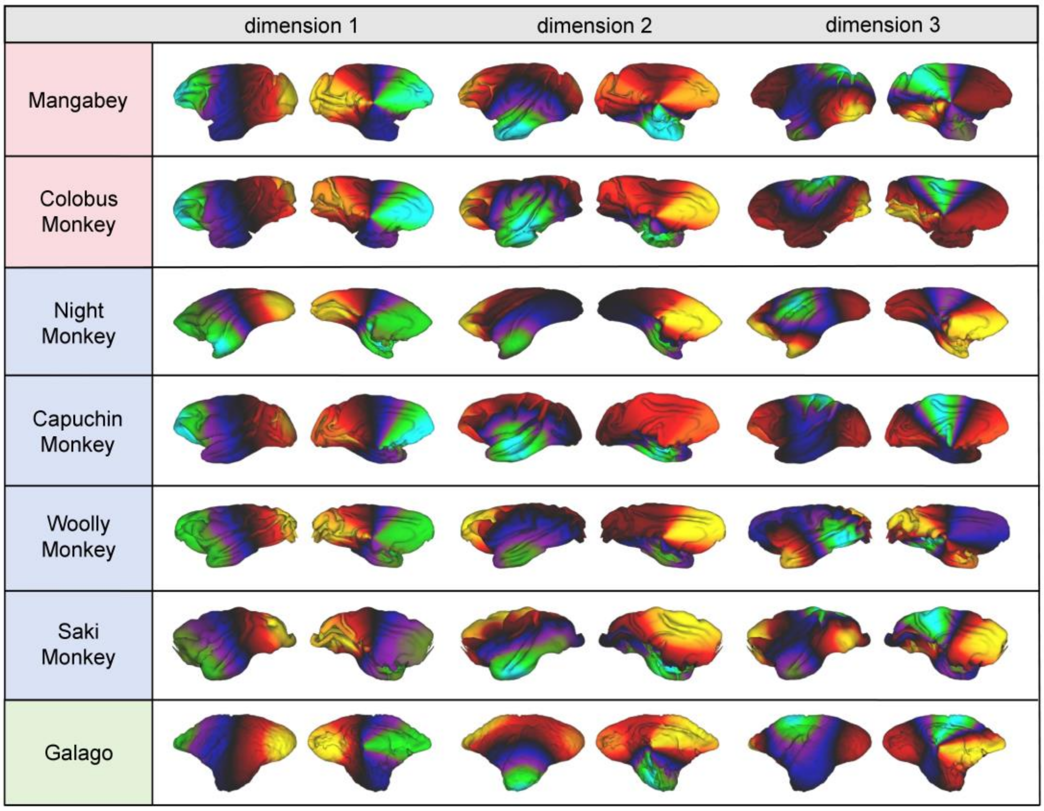


**Supplementary Figure 2.** Cross-species geometric gradients across extended primate species. The analysis was extended to eight additional primate species from different evolutionary branches, including two Old World monkeys (Mangabey and Colobus Monkey), four New World monkeys (Night Monkey, Woolly Monkey, Saki Monkey, and Capuchin Monkey), and one prosimian (Galago). The results reveal a conserved geometric organization across the primate lineage.

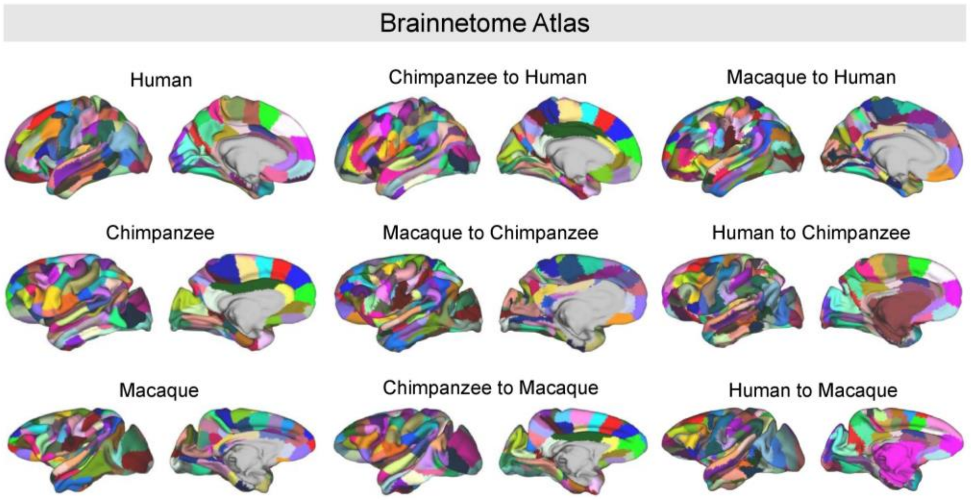


**Supplementary Figure 3.** Cross-species registration of Brainnetome Atlas. To evaluate the generalizability and applicability of the proposed cross-species registration framework, we applied the derived transformations to Brainnetome Atlas parcellations of human, chimpanzee, and macaque cortices.

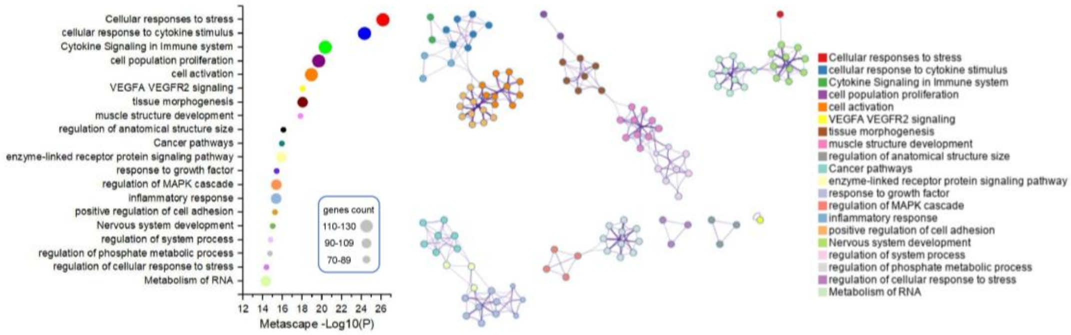


**Supplementary Figure 4.** Functional enrichment analysis of PLS1- genes.Gene Ontology (GO) enrichment results of PLS1− genes identified by PLS regression. (Left) Bubble plot of the top enriched biological processes from Metascape. Circle size indicates the number of genes per term, and color represents cluster identity. (Right) Network visualization of enriched GO terms, where each node represents an individual term and colors indicate functionally grouped clusters, revealing coherent functional modules associated with negatively weighted genes.
